# Clonal analysis of murine HSC self-renewal and differentiation in native hematopoiesis

**DOI:** 10.1101/2024.11.22.624575

**Authors:** Chenyu You, Zhen Zhang, Li Lin, Jianlong Sun

## Abstract

Hematopoietic stem cells (HSC) maintain lifelong hematopoiesis. However, in current clonal analyses with unbiased “barcoding” approaches, steady-state hematopoietic clones in young and middle-aged adults rarely have detectable HSCs, which precludes comprehensive interrogation of HSC clonal behaviors. In the current study, we used the previously described Sleeping Beauty transposon model to investigate HSC self-renewal and differentiation at a clonal level following a lifelong chase that significantly enriched HSC-derived clones. From seventeen mice, we detected over seventy thousand clones in native hematopoiesis that reflected the known HSC differentiation biases observed in transplantation. Our data indicated an intimate connection between megakaryocytic-restricted differentiation and HSC self-renewal expansion. By comparing the differentiation patterns of clones derived from transplanted HSCs, we further demonstrated the abilities of HSCs to preserve their cell fates towards self-renewal or multilineage differentiation. Unlike HSCs, clonal expansion in multipotent progenitors was associated with either a differentiation-active or differentiation-indolent state. Moreover, the clonal expansion events in the more differentiated stem and progenitor cells, but not the most primitive HSCs, drove clonal expansion in the megakaryocyte and myeloid cell lineages. Our study provided a comprehensive portrait of native hematopoiesis at a clonal level and revealed the general patterns in which HSCs maintained self-renewal and multi-lineage differentiation.

## Introduction

The hematolymphoid system consists of multiple types of terminally differentiated cells requiring continuous replenishment throughout life (Orkin & Zon, 2008). Maintenance of this regeneration program in adults relies on self-renewal and multilineage differentiation of hematopoietic stem and progenitor cells (HSPC) (Eaves, 2015). Studies using classical tools, such as single-cell transplantation (Ema et al., 2005; Takano, Ema, Sudo, & Nakauchi, 2004) and virus-mediated barcoding (Capel, Hawley, & Mintz, 1990; Jordan & Lemischka, 1990; Lemischka, Raulet, & Mulligan, 1986), demonstrate that these two hallmark characteristics can occur in single HSCs, laying the foundation for our current understanding of their basic biological properties. Since then, single-cell transplantation technology has revealed the heterogeneity of HSCs in terms of their self-renewal and differentiation potential (Carrelha et al., 2018; Dykstra et al., 2007; Ema et al., 2005; Muller-Sieburg, Cho, Karlsson, Huang, & Sieburg, 2004; Müller-Sieburg, Cho, Thoman, Adkins, & Sieburg, 2002; Sieburg et al., 2006; Yamamoto et al., 2013). The HSC compartment contains subsets showing balanced, myeloid-biased, lymphoid-biased, or megakaryocyte-restricted lineage outputs (Carrelha et al., 2018; Challen, Boles, Chambers, & Goodell, 2010; Dykstra et al., 2007; Morita, Ema, & Nakauchi, 2010; Muller-Sieburg et al., 2004; Sanjuan-Pla et al., 2013; Yamamoto et al., 2013). HSCs with long-term, intermediate-term, or short-term hematopoietic capacities are also recognized (Benveniste et al., 2010; Morita et al., 2010; Yang et al., 2005). These findings collectively indicate an HSC compartment with its constituents bearing different levels of self-renewal and differentiation potentials.

As myeloablation preconditioning is required for efficient donor engraftment in a transplantation assay, it has long been recognized that the hematopoietic behaviors of transplanted HSPCs can deviate from those displayed in unperturbed hematopoiesis (Hofer, Busch, Klapproth, & Rodewald, 2016; Konturek-Ciesla & Bryder, 2022; Pucella, Upadhaya, & Reizis, 2020). To address this problem, Cre recombinase-based lineage tracing approaches have been developed to label and track the self-renewal and differentiation behaviors of HSCs in their native microenvironment (Busch et al., 2015; Catherine et al., 2016; Chapple et al., 2018; Sawen et al., 2018). While the lineage tracing studies provide direct evidence of HSC differentiation, using a single-color fluorescent reporter in these studies prevents clonal analysis of HSC heterogeneity in situ. Although multi-color reporter systems have been used to perform lineage tracing with higher resolution (M. Ganuza et al., 2017; Miguel Ganuza et al., 2019; M. Ganuza et al., 2022; Kucinski et al., 2024; Liu et al., 2024; Yu et al., 2016), the number of HSCs present in adults surpasses the number of distinct colors offered by the reporter system, making unique tagging of single HSCs a challenging task. Along with the Cre-based lineage tracing studies, a series of tools to create de novo “barcode” in the genome are developed (Bowling et al., 2020; Feng et al., 2022; L. Li et al., 2023; Pei et al., 2017; Pei et al., 2020; J. Sun et al., 2014). These DNA-based barcodes are derived from random transposon genome insertion sites (Feng et al., 2022; J. Sun et al., 2014), random recombination among tandem *loxP* sites (Pei et al., 2017; Pei et al., 2020), and random indel sequences following the repair of CRISPR-mediated double-strand break (Bowling et al., 2020; L. Li et al., 2023). As these methods offer large numbers of unique barcodes far exceeding the number of HSPCs present in these murine models, the individual HSPCs in these models can be labeled distinctively for downstream clonal analysis (Bowling et al., 2020; Feng et al., 2022; L. Li et al., 2023; J. Sun et al., 2014). The DNA-based barcode systems are ideal models for analyzing HSC heterogeneity. However, most current barcode tools label HSPCs in a non-cell type-specific manner. As a result, the identification of HSC-derived clones relies exclusively on detecting the common barcodes in HSCs and their downstream progenies (Bowling et al., 2020; L. Li et al., 2023; J. Sun et al., 2014). While such HSC-rooted clones are common during early embryonic development or in stress hematopoiesis (Bowling et al., 2020; L. Li et al., 2023; Pei et al., 2017), they are rare under the steady-state condition in young and middle-aged adults (Bowling et al., 2020; L. Li et al., 2023; Pei et al., 2017; J. Sun et al., 2014). This problem is potentially associated with the limited efficiency of barcode detection and cell sampling, making clonal identification technically challenging for the rare differentiating HSCs without sufficient self-renewal expansion.

The HSC compartment expands significantly over time, potentially caused by the HSCs’ increased tendency to self-renewal during aging (Itokawa et al., 2022; Rossi et al., 2005; D. Sun et al., 2014). Moreover, lineage tracing studies demonstrate gradually accumulated contribution from the adult HSCs and washout of progeny from multipotent progenitors (Busch et al., 2015; Catherine et al., 2016; Chapple et al., 2018; Patel et al., 2022; Sawen et al., 2018). Therefore, we reasoned that increasing the chasing period would enrich HSC-derived clones, and more of these clones would contain HSCs due to increased HSC self-renewal expansion. With this consideration, we analyzed seventeen aged mice carrying the Sleeping Beauty transposon labeling system (M2/HSB/Tn) to detect HSC-derived clones. These mice were induced for transposon labeling at young ages and analyzed approximately two years later to detect transposon chromosomal integration sites (Tn tags) in HSPCs and mature cell types. A comparison of these detected tags in the different HSPC populations and downstream cells allowed the retrospective analysis of HSC differentiation histories. Our results comprehensively overview HSC clonal behaviors in steady-state hematopoiesis and reveal the major patterns of HSC self-renewal and differentiation.

## Results

### Examination of hematopoietic clonal behaviors with Sleeping Beauty transposon labeling

To understand the clonal behaviors of HSPCs, we used the previously described Sleeping Beauty transposon (Tn) system to label cells uniquely (J. Sun et al., 2014). Seventeen mice with the Sleeping Beauty lineage tracing system from seven cohorts were fed with doxycycline (Dox) for 4 weeks beginning at a young age (6-10 weeks). Tn tags in labeled (DsRed^+^) HSC with the high (EPCR-high HSC: E^hi^ HSC) or low (EPCR-low HSC, EPCR-negative HSC: E^lo^/E^neg^ HSC) self-renewal ability (Balazs, Fabian, Esmon, & Mulligan, 2006; Schulte et al., 2015; Umemoto et al., 2022), hematopoietic progenitor cells (HPC-1 and HPC-2), megakaryocyte progenitors (MkP), myeloid progenitors (MyP), granulocytes (Gr), monocytes (Mono), B cells, and T cells FACS-purified from bone marrow of femurs, tibias, and spines were analyzed approximately 100 weeks (90-120 weeks) after Dox withdrawal (Figure 1A, Figure 1—figure supplement 1A, B, Supplementary file 1). Approximately 20-40% of the cells were labeled in these aged mice, and spontaneous Tn tag labeling, as revealed by DsRed^+^ cells in the absence of Dox induction, occurred at a minimal level except for the T cells (Figure 1—figure supplement 1C). Tn tags in FACS-purified DsRed^+^ cells were detected by TRACE (transposase-assisted capture of transposable elements) (Patel et al., 2022), a method that incorporated unique molecular identifiers (UMIs) into library preparation to measure Tn tag abundance semi-quantitatively (Figure 1—figure supplement 2A, B).

**Figure 1:**
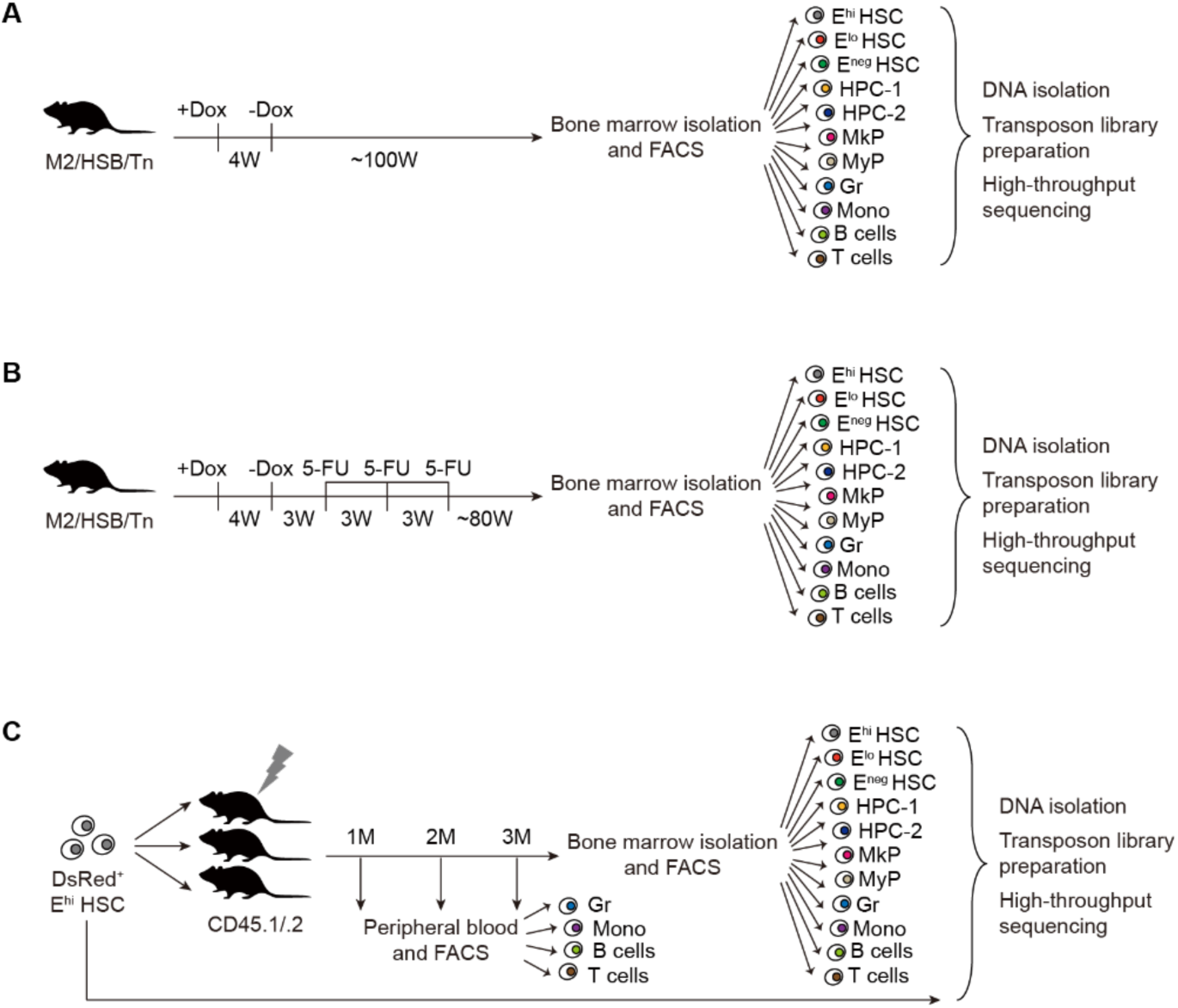
Clonal analysis of murine hematopoiesis. A: Experimental outline for lineage tracing of unperturbed hematopoiesis. M2/HSB/Tn mice were labeled with Dox for four weeks. The DsRed^+^ fractions of the indicated cell populations were isolated from bone marrow approximately a hundred weeks after the removal of Dox for Tn tag library preparation and sequencing. B: Experimental outline for lineage tracing of hematopoiesis following 5-FU treatment. Three weeks after Dox withdrawal, the M2/HSB/Tn mice received three rounds of 5-FU injections at a 3-week interval. The indicated DsRed^+^ cell populations were isolated from bone marrow approximately eighty weeks after the last 5-FU infusion. C: Experimental outline for comparative analysis of E^hi^ HSC clonal behaviors in native hematopoiesis and after bone marrow transplantation. One-fourth of the isolated donor DsRed^+^ E^hi^ HSCs were used for Tn tag detection. The rest cells were transplanted into lethally irradiated mice. The DsRed^+^ fractions of the indicated cell populations were isolated monthly from peripheral blood. The recipient mice were sacrificed three months after transplantation to analyze Tn tags in bone marrow cell populations.

To elucidate the clonal behaviors of HSCs with a further increased proliferation history, we treated five of these mice three weeks after Dox induction with three rounds of 5-Fluorouracil (5-FU), a treatment previously used to model accelerated aging (Beerman et al., 2013) (Figure 1B). These mice were sacrificed for Tn tag analysis approximately 80 weeks after the last 5-FU treatment due to their poorer health conditions than the unperturbed mice. At the time of Tn tag analysis, we transplanted three-quarters of the Tn-labeled E^hi^ HSCs from two naturally aged and two 5-FU-treated mice into syngeneic CD45.1^+^ mice to reveal the impacts of transplantation on HSC clonal behaviors (Figure 1C).

### Diverse differentiation patterns are detected in unperturbed hematopoiesis

A total of 73,452 clones were detected from the seventeen mice with and without 5-FU perturbation (Figure 2A, Figure 2—figure supplement 1, Supplementary file 2). Approximately 18.5% of the clones contained more than two cell types, and the rest contained only one (Figure 2A). In the following analysis, these clones were referred to as “multi-cell type” and “unique” clones, respectively.

**Figure 2:**
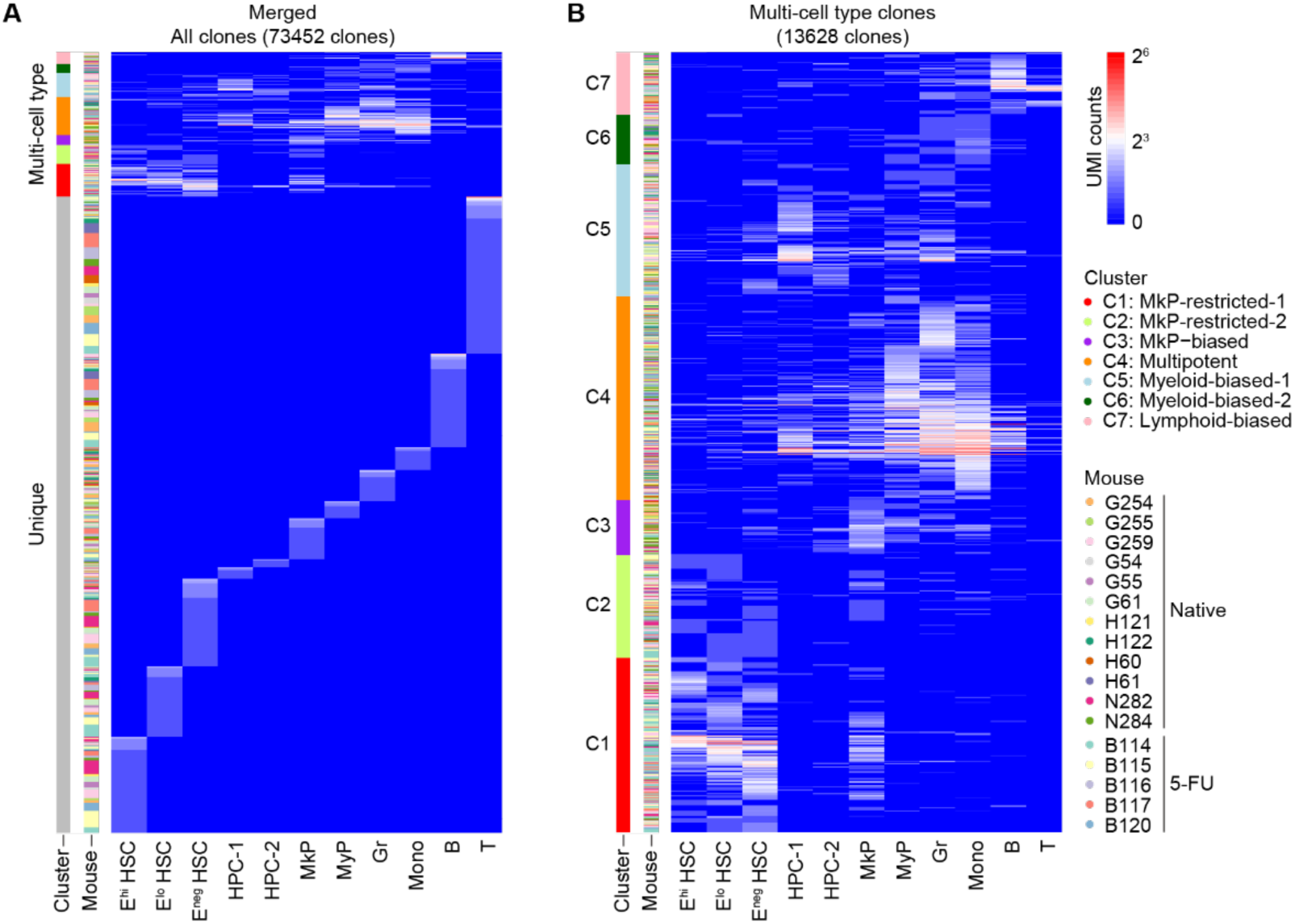
Differentiation patterns of hematopoiesis in unperturbed and 5-FU-treated mice. A: Heatmap depicting the alignment of Tn tags from all mice examined, with each row representing a unique Tn tag and each column representing the cell types. B: Heatmap showing the result of Louvain clustering on multi-cell type clones. Tn tags are colored by log2-transformed UMI counts, which reflect their abundance. Tn tags used in the plots are pooled from mice assayed in the unperturbed condition (n = 12) and after 5-FU treatment (n = 5).

As the Tn tag analysis was done in cells harvested after a lifelong chase, previous lineage tracing studies using HSC-specific Cre drivers suggest that most of these clones present after such a long chase period, especially those containing the short-lived myeloid cells, were derived from HSCs (Busch et al., 2015). In addition, 3206 multi-cell type clones contained the most primitive E^hi^ HSC. These E^hi^ HSC-containing clones provided direct experiment evidence to evaluate the relationship between HSC self-renewal and differentiation.

The different cell types in the multi-cell type clones represented the footprints of differentiation events in these clones, which helped reveal their differentiation patterns. However, like most other types of barcode analysis, the sampling issue and limited sensitivity of Tn tag detection prevented reliable calls if particular cell types were present or absent in these clones. On the other hand, these ambiguities were expected to occur randomly, and the analysis of a large number of clones detected in the current study with advanced statistical tools would help to reveal the general differentiation patterns in native hematopoiesis. With these considerations, we focused on the multi-cell type clones to determine the patterns of hematopoietic differentiation using Louvain clustering based on individual clones’ UMI counts in their detected cell types. This analysis revealed seven clusters of clones that differed in their compositions of differentiated cells: C1, MkP-restricted-1; C2, MkP-restricted-2; C3, MkP-biased; C4, multipotent; C5, myeloid-biased-1; C6, myeloid-biased-2; C7, lymphoid-biased (Figure 2B). These clone types recapitulated the major differentiation patterns of transplanted HSCs (Carrelha et al., 2018; Challen et al., 2010; Dykstra et al., 2007; Morita et al., 2010; Muller-Sieburg et al., 2004; Sanjuan-Pla et al., 2013; Yamamoto et al., 2013), indicating that the HSCs displayed similar heterogeneity in their differentiation behaviors under the native and transplantation conditions. A manual examination of the multipotent C4 clones revealed over half of the clones being myeloid-biased (Figure 2B). Therefore, myeloid-biased clones were among the most prevalent clone types in aged mice, which was highly expected given the myeloid-biased differentiation pattern observed during aging (Rossi et al., 2005; Sudo, Ema, Morita, & Nakauchi, 2000; Yamamoto et al., 2018). The clones detected in the 5-FU-treated mice were not enriched in particular clone types (Figure 2B), indicating negligible effects of increased HSC proliferation histories on major hematopoietic differentiation patterns.

### MkP-restricted differentiation pattern is associated with HSC expansion

Although most clones detected after the lifelong chase, except for the long-lived lymphoid clones, were likely HSC-derived, the HSCs were detected in variable frequencies in these clones. Compared to the rest clone types, many more clones in the MkP-restricted C1 and C2 clone clusters contained HSCs (Figure 3A). Moreover, the average UMI counts of the E^hi^ HSCs, E^lo^ HSCs, and E^neg^ HSCs were higher in the C1 MkP-restricted clones of unperturbed mice (Figure 3B). This result indicates that the HSCs in the C1 MkP-restricted clones experience more extensive self-renewal expansion than HSCs in other clone types.

**Figure 3:**
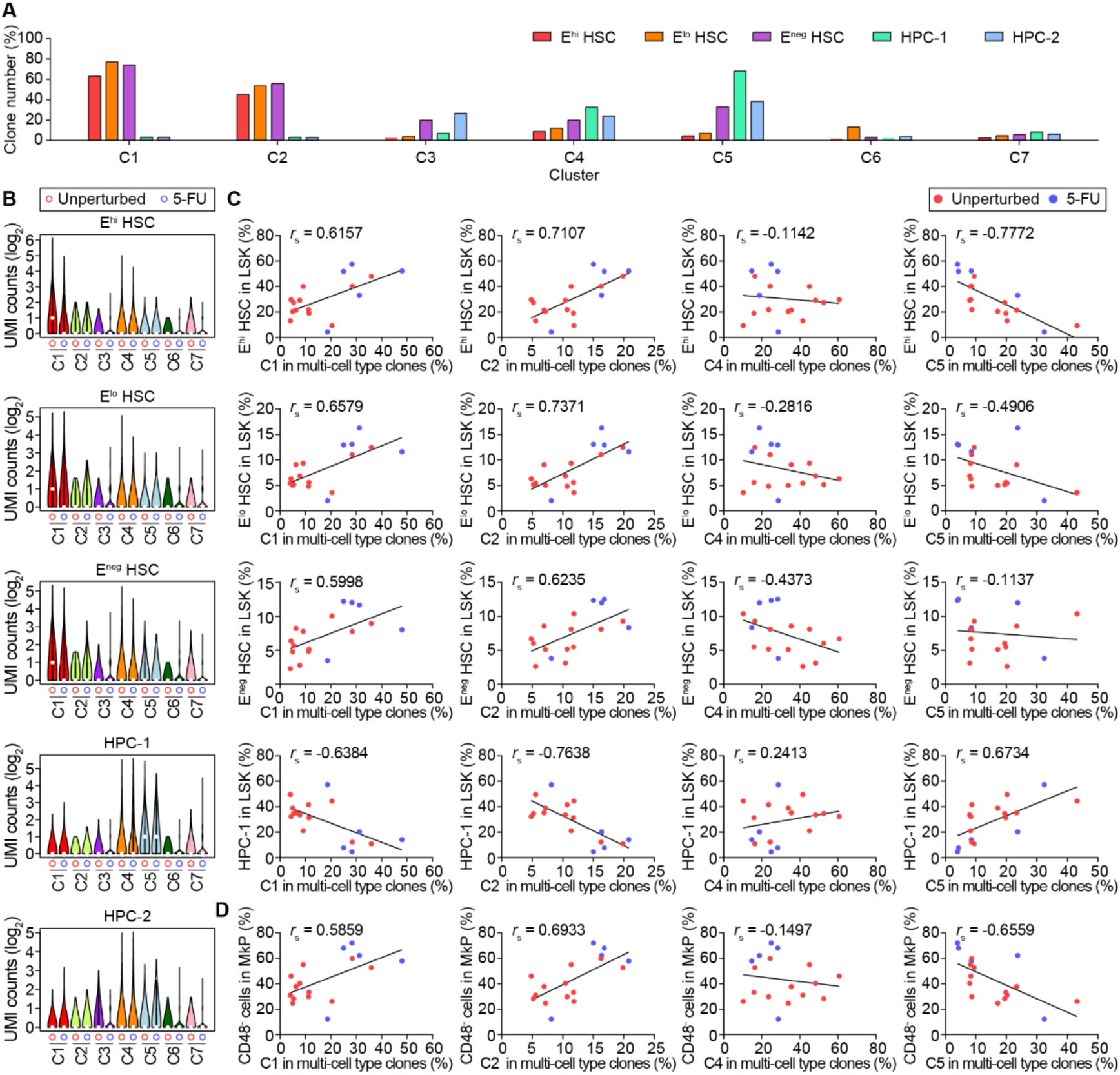
The correlation between HSC expansion and MkP-restricted differentiation. A: The percentage of clones in each cluster that contained the indicated cell types. B: Violin plots showing the Tn tag UMI counts of the indicated cell types in each clone cluster. Tn tags are pooled from unperturbed mice (red circle, n = 12) and 5-FU-treated mice (blue circle, n = 5). C: Correlations between the frequencies of the indicated multipotent hematopoietic cell populations in Lin^-^ cKit^+^ Sca1^+^ (LSK) cells and the frequencies of the indicated clone cluster in multi-cell type clones. Each dot represents data from a single mouse (unperturbed: red, n = 12; 5-FU: blue, n = 5). “*r*_s_” depicts the Spearman correlation coefficient. D: Correlations between the frequencies of CD48^-^ cells in MkP and the frequencies of the indicated clone cluster in multi-cell type clones. Each dot represents data from a single mouse (unperturbed: red, n = 12; 5-FU: blue, n = 5). “*r*_s_” depicts the Spearman correlation coefficient.

Further supporting the correlation between HSC self-renewal expansion and MkP-restricted differentiation, we found that the individual mice’s C1 clone frequency was positively correlated with their E^hi^/E^lo^/E^neg^ HSC frequencies (Figure 3C). Such a correlation was also observed for the C2 MkP-restricted clones but not for the C4/C5 clones (Figure 3C) and the C3/C6/C7 clones (data not shown). This positive correlation was most evident in four of the five 5-FU-treated mice, whose HSC frequencies and C1/C2 clone abundance were higher than those from the unperturbed mice (Figure 3C, Figure 3—figure supplement 1A). In addition to the correlation with HSC frequencies, mice with more C1 or C2 clones also displayed a higher frequency of the CD48^-^ MkP cells (Figure 3D, Figure 3—figure supplement 1B). This observation supports a recent lineage tracing study’s conclusion that the CD48^-^ MkPs are directly derived from HSCs (J.-J. Li et al., 2024).

Collectively, these findings suggest that HSCs with an MkP-restricted differentiation pattern have a higher chance of undergoing self-renewal expansion than those undergoing multilineage or myeloid-biased differentiation.

### HPC-1 contains a differentiation-active and a differentiation-indolent subgroup

In addition to HSCs, we observed exceptional expansion of HPC-1, as revealed by their much higher average UMI counts, in the C5 clones than other clone types (Figure 3B). Such a clone type-specific expansion did not occur to the HPC-2 cells (Figure 3B). Correspondingly, we observed a positive correlation between C5 clone frequency and HPC-1 in the LSK cells (Figure 3C). While the HPC-1 cells were also frequently detected in the C4 clones (Figure 3A), we did not observe a similar correlation between C4 clone abundance and HPC-1 frequency (Figure 3C). These observations imply that the HPC-1 in the C4 and C5 clones may reside in different self-renewal states, leading to different levels of clonal expansion in the HPC-1 pool.

Indeed, a close inspection of the C4 and C5 clones showed much fewer MyP cells, granulocytes, and monocytes in C5 clones than in C4 clones (Figure 2B). This phenomenon led us to speculate that the HPC-1 cells might be heterogeneous in their differentiation behaviors: those in the C4 clones were actively producing downstream cells, whereas the others in the C5 clones were in a self-renewal-biased and differentiation-indolent state. To test this hypothesis, we employed the Gaussian-Mixture-Model (GMM) implemented in R package mclust (Scrucca, Fraley, Murphy, & Adrian E., 2023) to re-cluster the clones in C4 and C5 based on the log2 ratio of their UMI counts in MyP, Gr, or Mono against HPC-1 (UMI_MyP_ / UMI_HPC-1_, UMI_Gr_ / UMI_HPC-1_, or UMI_Mono_ / UMI_HPC-1_). We assumed that a high UMI_MyP/Gr/Mono_ / UMI_HPC-1_ ratio reflected an active differentiation state and a low ratio reflected an indolent differentiation state of the HPC-1 clones. If GMM suggested one component model (i.e. the HPC-1 cells were in the same differentiation state), the variance of the log2 ratios was mainly derived from random noise. On the contrary, if GMM suggested a two-component model, the variance of ratios could be better explained by the existence of two different differentiation states. We fitted the model to log2 ratios calculated from the three pairs (MyP/HPC-1, Gr/HPC-1, and Mono/HPC-1) in all C4 and C5 clones. For all three pairs, the GMM suggested that models with more components were better than a one-component model based on BIC (Bayesian Information Criterion) (Figure 4A), suggesting that diverse differentiation states were indeed present in HPC-1. For simplicity, we only considered a two-component model in the following analyses. We compared the observed distribution of the log2 UMI_MyP/Gr/Mono_ / UMI_HPC-1_ ratios against the predicted distribution generated based on the one-component or two-component model (Figure 4B). In all three cases, a visual inspection confirmed that the two-component model better fitted the experimental data than the one-component model (Figure 4B).

**Figure 4:**
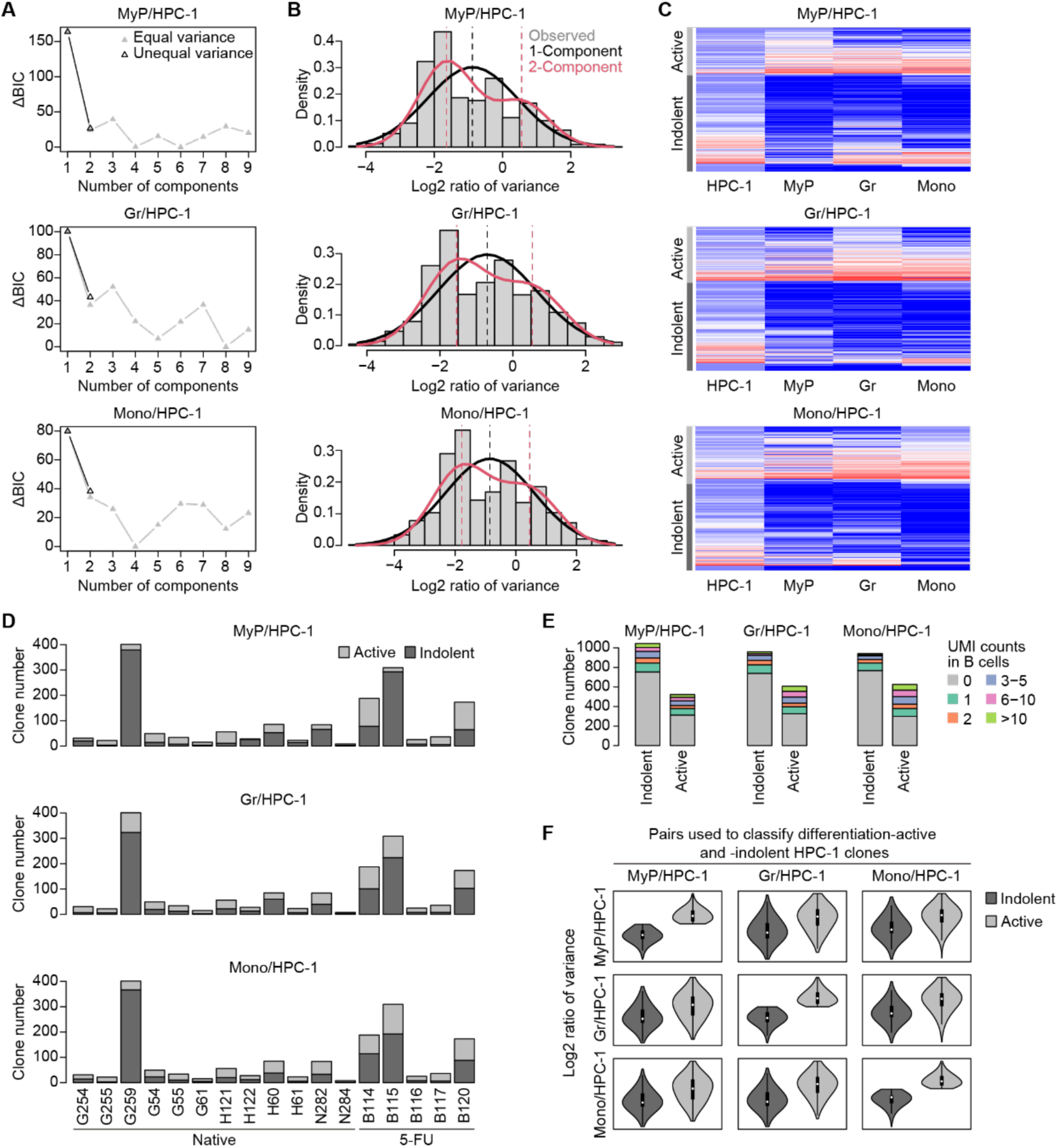
The heterogeneous differentiation states of HPC-1. A: The Δ Bayesian information criterion (ΔBIC) demonstrates the relative difference between the best model that explains the log2 ratios of Tn tag UMI counts in downstream differentiated cell populations and upstream HPC-1 (down/up pairs: MyP/HPC-1, Gr/HPC-1, or Mono/HPC1), and Gaussian mixture models (GMM) with varying numbers of components. The results from models that assume each component has equal variance (gray) and unequal variance (black) are both shown. B: The observed distributions of the log2 ratios of Tn tag UMI counts in the three down/up pairs (gray bar) are shown alongside the distributions predicted by the one-(black curve) or two-component (red curve) Gaussian mixture models with minimal ΔBIC. The dashed lines illustrate the mean values predicted by the one-(black) or two-component (red) Gaussian mixture models. C: Heatmaps showing the classification results of differentiation-active and -indolent clones defined based on their log2 UMI count ratios in the three down/up pairs using a two-component Gaussian mixture model with minimal ΔBIC. Tn tags are colored by log2-transformed UMI counts. D: The number of differentiation-active and -indolent clones defined based on their log2 UMI count ratios of the three down/up pairs in each mouse. E: The number of clones with indicated B-cell UMI counts in differentiation-active and - indolent HPC-1 clones. G: Violin plots showing the distribution of the log2 ratios of Tn tag UMI counts in the three down/up pairs in differentiation-active and -indolent HPC-1 clones classified by the individual down/up pairs. Except for Figure 5D, data used in this figure is pooled from mice assayed in the unperturbed state (n = 12) and after 5-FU-induced regeneration (n = 5).

Based on the two-component model, the C4 and C5 clones were re-classified into differentiation-active and differentiation-indolent groups according to their log2 UMI_MyP/Gr/Mono_ / UMI_HPC-1_ ratio (Figure 4C). We found that the two groups of clones could be simultaneously found in most mice (Figure 4D), indicating that heterogeneity in differentiation states was a common phenomenon in all mice examined. The HPC-1 could be divided into two sub-groups based on the expression of Flt3, a maker indicating lymphoid differentiation priming (Adolfsson et al., 2005). Therefore, we asked whether the different differentiation states of the HPC-1 clones were associated with a lymphoid differentiation bias. We found that both groups had substantial B-cell-producing clones (Figure 4E). As expected, the differentiation-active clones had a larger fraction of B-cell-producing clones than the differentiation-indolent clones (Figure 4E). However, most of these B-cell-producing clones also contained myeloid cells (Figure 2B), indicating that the differentiation-active clones were not lymphoid-biased but, instead, produced more myeloid and lymphoid cells simultaneously.

Finally, we asked whether the inferred differentiation-indolent and -active clones identified based on ratios of the MyP/HPC-1, Gr/HPC-1, or Mo/HPC-1 pairs were consistent. Although the digital classifications inferred from three pairs were not matched well by the Adjusted Rand Index (MyP vs. Gr, ARI = 0.21; MyP vs. Mono, ARI = 0.23; Gr vs. Mono, ARI = 0.32), the differentiation-indolent clones inferred by any of the three pairs tended to have a lower ability to produce all downstream cells (Figure 4F). This result indicates concurrent lineage output in MyP and the two myeloid cell lineages if the HPC-1 clones are in a differentiation-active state.

Next, we examined whether such heterogeneity of differentiation ability also existed in MyP. We performed the same analyses on the two progenitor-downstream pairs (Gr/MyP and Mono/MyP). GMM suggested a two-component model was better than a one-component model for Gr/MyP. However, the ΔBIC was smaller than the cases in HPC-1, indicating that the heterogeneity was less significant (Figure 4—figure supplement 1A). For Mono/MyP, the two-component model did not fit the data better than the one-component model (Figure 4—figure supplement 1B). We further compared the predicted and observed density of the log2 ratio. Consistent with the results of the statistical test by BIC, the two-component model did not show significant improvement compared to the one-component model (Figure 4—figure supplement 1B). Moreover, we found that differentiation-active clones classified by the two-component model had smaller clonal sizes (Figure 4—figure supplement 1C), suggesting that their increased log2 ratio might be derived from the higher noise associated with these clones’ low MyP UMI counts. In conclusion, the data did not strongly support the heterogeneity of differentiation ability in MyP.

Collectively, the above analysis revealed two types of HPC-1 subsets residing in a differentiation-active or -indolent state. The frequency of differentiation-indolent HPC-1 clones, found predominantly in C5 clones and, to a lesser degree, in C4 clones, may contribute to HPC-1 expansion.

### Clonal expansion is prevalent in hematopoietic cells of aged mice

After analyzing the connection between HSPC self-renewal and distinct differentiation patterns, we next focused on clonal hematopoiesis, a phenomenon prevalent during aging in humans and mice (Miguel Ganuza et al., 2019; Jaiswal, 2020; Mitchell et al., 2022; van Zeventer et al., 2021). Consistent with the literature, we observed Tn tags with large UMI counts, indicating clonal expansion, in most cell types examined (Figure 2A, Figure 2—figure supplement 1). To evaluate the levels of clonal expansion, we calculated each cell population’s Pielou’s index, a robust and sensitive indicator for clonal dominance, with the adjustment for the sample size effect (Corre & Galy, 2023). Pielou’s indexes of all cell populations from aged mice were significantly lower than the freshly induced granulocytes (Figure 5A), indicating a widespread increase in clonal dominance over the long chasing period. We found that the 5-FU treatment had a limited impact on Pielou’s index of E^hi^/E^lo^/E^neg^ HSCs and MkPs (Figure 5A), indicating a relatively even expansion of HSC clones after 5-FU-induced hematopoietic regeneration. Moreover, MkP could be produced directly by HSCs (Carrelha et al., 2018; J.-J. Li et al., 2024; Poscablo et al., 2024), potentially preventing their loss of clonal complexity. In contrast to HSCs and MkP, 5-FU significantly reduced the clonal complexity in other progenitors and differentiated cell types, except for the T-cells. This outcome was consistent with the fact that the proliferating progenitors were depleted by 5-FU treatment and replenished by the progeny of much rarer HSCs in 5-FU-induced hematopoietic regeneration (Busch et al., 2015; Lerner & Harrison, 1990).

**Figure 5:**
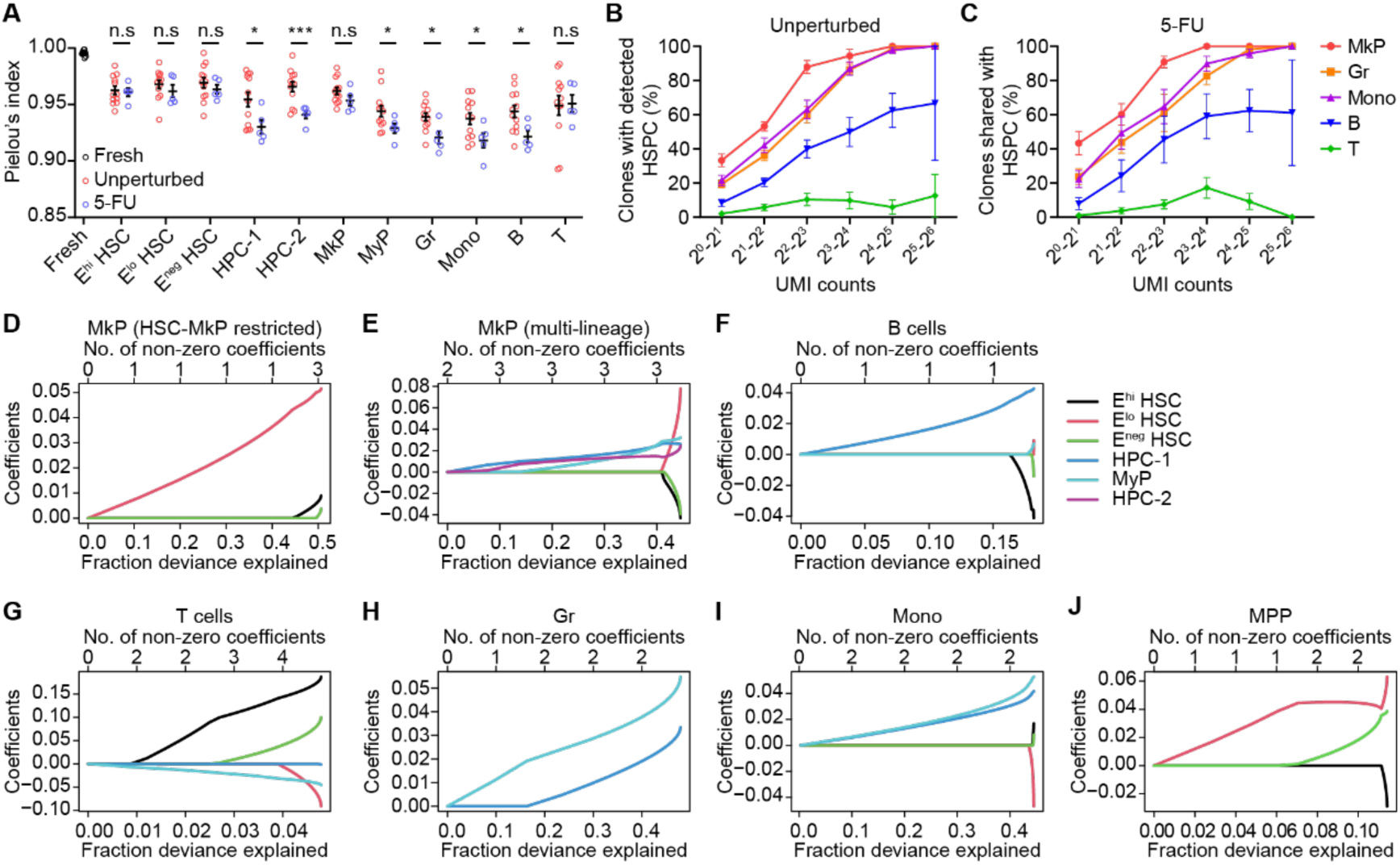
The cellular origins for clonal expansion in mature lineage blood cells. A: Pielou’s index measures the evenness of UMI counts of Tn tags detected in the freshly induced granulocytes (Fresh) and all assayed bone marrow cell populations. Each circle represents data from a technical repeat (Fresh, n = 12) or a single mouse (unperturbed: red, n = 12; 5-FU: blue, n = 5). Bars represent means with SEM. Two-tailed unpaired Student’s t-tests are used to compare data from two experimental groups. ns, not significant, *p < 0.05, ***p < 0.001. B and C: Frequency of clones detected in downstream mature lineages (MkP, Gr, Mono, B cells, T cells) with different UMI counts whose Tn tags also detected in multipotent hematopoietic stem and progenitor cells (HSPC: E^hi/lo/neg^ HSC, HPC-1, HPC-2) (B: unperturbed, n = 12; C: 5-FU, n = 5). Bars represent means with SEM. D-J: The results of ElasticNet illustrates the linear relationships of UMI counts of Tn tags detected in downstream cell populations (MkP: C1/C2 (D), C3-C7 (E); B cells (F); T cells (G); Gr (H); Mono (I); and HPC-1 (J)) with their upstream progenitors (HSC: E^hi/lo/neg^ HSC (D, J); E^hi/lo/neg^ HSC, HPC-1, HPC-2, MyP (E); E^hi/lo/neg^ HSC, HPC-1, MyP (F-I)). The x axes represent a series of models with different penalty settings. The fraction of deviance explained and the number of non-zero coefficients in the corresponding models are shown separately in the lower and upper bound. The y axes show the coefficients (*a_i_*) estimated in these models. Data in this plot is pooled from mice assayed in the unperturbed state (n = 12) and after 5-FU-induced regeneration (n = 5).

### Clonal expansion in different blood cell lineages has diverse cellular origins

In unperturbed and 5-FU treated mice, we observed a positive correlation between each clone’s expansion (UMI counts) in their differentiated cell types and their chance of detecting the HSPCs (Figure 5B, C), implying potential roles of upstream HSPCs in driving clonal expansion in the downstream cell types. To formally test this hypothesis, we asked whether the clonal expansion in the individual differentiated cell populations was inherited from one of their upstream progenitors or derived from their own self-renewal expansion. We hypothesized that if the clonal expansion in one cell population was inherited from its progenitors, its clonal size variations could be explained mainly by the clonal size variations in its progenitors. Thus, we used ElasticNet to investigate the linear relationships of UMI counts in one cell population and all of its possible upstream progenitors (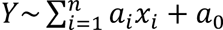, where *Y* corresponds to the number of UMIs in examined populations, *x_i_* corresponds to the number of UMIs of its *i*th progenitor, *a_i_* corresponds to the coefficient of *x_i_* and *a_0_* corresponds to the intercept). The ElasticNet was a generalized linear model via penalized maximum likelihood, which uses the L1-or L2-norm of *a_i_* to control the number of *x_i_* (the model complexity) to explain the response *Y* (Friedman, Hastie, & Tibshirani, 2010; Simon, Friedman, Hastie, & Tibshirani, 2011).

We first investigated the linear relationships between MkP and its progenitors. As the MkPs were found in the MkP-restricted clusters C1/C2 and non-restricted clusters (C3-C7), we analyzed MkP clone origins in these clusters separately. In MkP-restricted C1/C2 clones, the model with one non-zero coefficient (E^lo^ HSC) could explain up to 44% UMI count deviance in MkP. In comparison, the full model with three non-zero coefficients, or all three HSC types regulating MkP expansion, could only explain 51% deviance in MkP, and E^lo^ HSC also had the highest coefficient (Figure 5D). As adding more complexity only marginally improved the prediction power, this result strongly suggested that the clonal expansion of MkPs in C1/C2 was inherited from E^lo^ HSC. In contrast to the C1/C2 clones, the sizes of MkP clones in other MkP-producing clones (C3-C7) could be explained by the clonal sizes of HPC-1, HPC-2, and MyP with similar coefficients (Figure 5E), suggesting the clonal expansion in these MkPs might be inherited from multiple progenitors.

We next investigate the clonal expansion in myeloid and lymphoid cells. From the results of ElasticNet, the UMI count deviance in B and T cells could not be well explained by their progenitors (Figure 5F, G), indicating that the clonal expansions of B and T cells were mainly derived from self-expansion but not inherited from their progenitors. In contrast, the sizes of each clone in Gr and Mono could be explained by that in HPC-1 and MyP (Figure 5H, I), suggesting that clonal expansion in these myeloid populations was mainly inherited from the HPC-1 and MyP stage. On the other hand, only around 10% of the variance in HPC-1 expansion was explained by the upstream HSC cell types (Figure 5J), indicating that clonal expansion in the myeloid cell populations was determined at the HPC-1 but not the HSC levels.

Together, these results indicate that clonal expansion of a subset of the lineage-restricted MkP inherited from the E^lo^ HSC, whereas HPC-1 and MyP mostly drove the clonal expansion events in the multipotent MkP and myeloid cell compartments. In contrast to MkP and myeloid cells, clonal expansion in the lymphocytes occurred primarily in the lymphoid lineages.

### Clonal behaviors of transplanted HSCs

Finally, we analyzed the clonal behaviors of E^hi^ HSCs after bone marrow transplantation. The four donor mice, two with prior 5-FU treatment and two with no perturbation contained all types of unique and multi-cell type clones (Figure 6—figure supplement 1A, B), indicating that the clones in these four donor mice represented all the observed clone types. By comparing the donor and recipient clones, we identified 542 donor clones in the recipients (Figure 6A). Based on these clones, we calculated the transplantation survival probability of each donor clone type. The survival probability of the individual donor clone types varied significantly, with the C1 clones, the clone type containing most of the expanded HSC clones, showing the highest survival probability (Figure 6B). A similar survival probability among clones from 5-FU-treated mice was observed (Figure 6C). Interestingly, despite some of the donor clone types, such as the MkP unique clones, the C3 MkP-biased clones, and the C4/C5/C6 multi-cell type clones, did not contain many HSCs, they were recovered and contributed to a significant portion of the 542 recurrent clones (Figure 6B, C).

**Figure 6:**
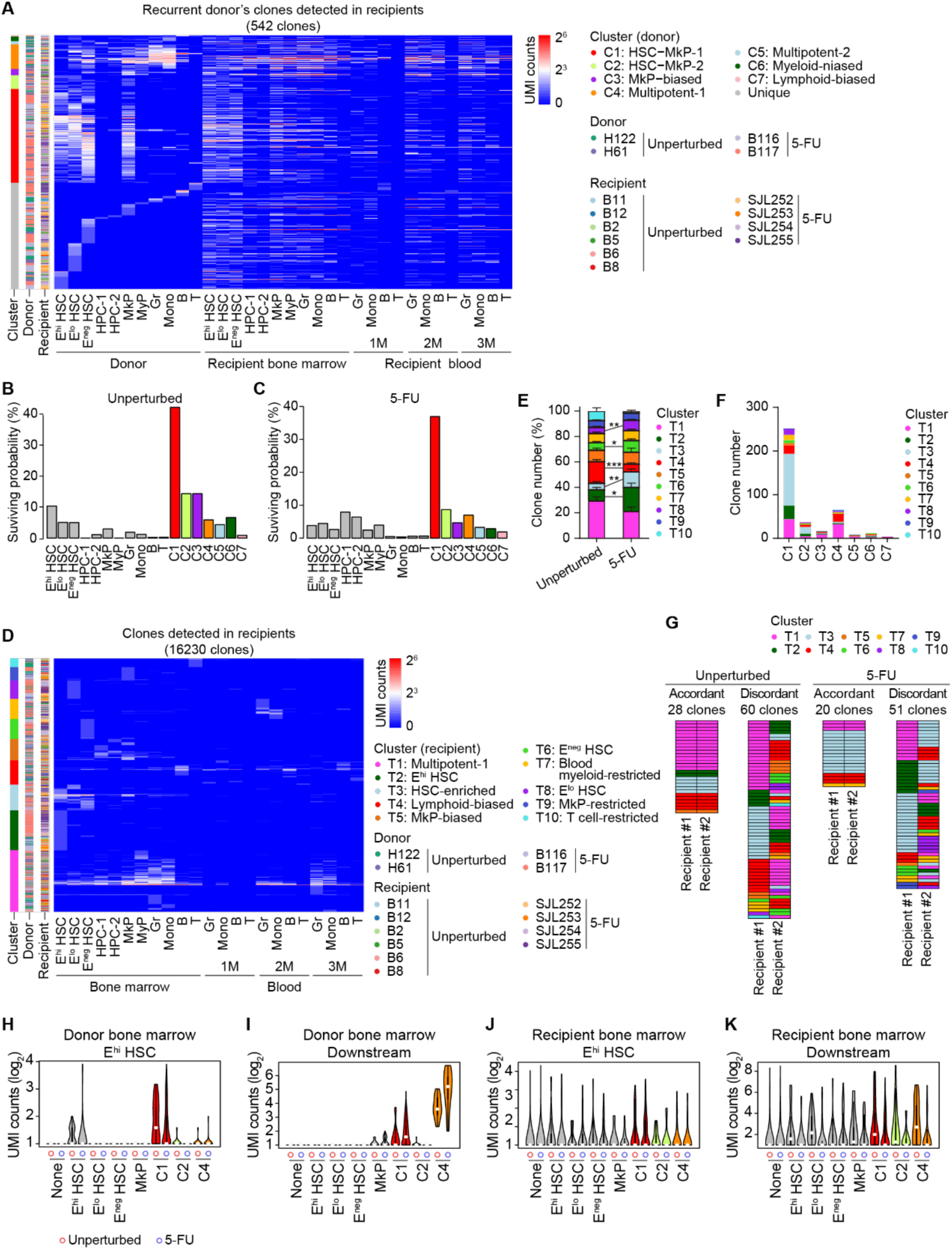
Clonal analysis of hematopoiesis after transplantation. A: The heatmap showing donor clones detected in recipient mice after transplantation. Tn tags are colored by log2-transformed UMI counts. The transposon tags used in the plot are pooled from donor mice used for transplantation (n = 2 for each experimental group). B and C: Survival probability of different donor clone types measured by the fraction of donor clones detected in transplant recipients in all clones in the corresponding clone type. B: donor clones from unperturbed mice; C: donor clones from 5-FU treated mice. D: The heatmap showing the result of Louvain clustering on clones detected in the recipients’ bone marrow and blood cell populations. Tn tags are colored by log2-transformed UMI counts. The Tn tags used in the plot are pooled from recipient mice transplanted with donor E^hi^ HSCs from the unperturbed group (n = 6) and the 5-FU group (n = 4). E: The average frequency of the individual recipient clone types in mice receiving donor E^hi^ HSCs from unperturbed (n=6) and 5-FU-treated mice (n=4). Bars represent means with SEM. Two-tailed unpaired Student’s t-tests are used to compare data from two experimental groups. *p < 0.05, **p < 0.01, ***p < 0.001, ***p < 0.001. F: The number of recipient clone types generated by the seven types of donor multi-cell type clones. Clones from the unperturbed and 5-FU-treated donor mice are pooled for the calculation. G: Alignment of recipient clones detected in two recipients. The rows represent clones, and the columns represent recipient mice. Clones are colored according to their cluster types in the recipient mice from which they are derived. H-K: Violin plots showing the UMI counts of the different types of donor clones in the donor E^hi^ HSC (H), the donor mature cell lineages (Gr, Mono, B-cells, T-cells) (I), the recipient E^hi^ HSC (J), and the recipient mature cell lineages (Gr, Mono, B-cells, T-cells) (K). Clones from unperturbed and 5-FU-treated donors and recipients are separately pooled for the analysis.

We compared the differentiation patterns of these surviving clones in donors and recipients (Figure 6D). For this purpose, we performed similar Louvain clustering on all the clones detected in the recipients, which revealed ten clusters of clones (Figure 6D): T1, multipotent-1; T2, E^hi^ HSC-restricted; T3, HSC-enriched; T4, lymphoid-biased; T5, MkP-biased; T6, E^neg^ HSC-restricted; T7, blood myeloid-restricted; T8, E^lo^ HSC-restricted; T9, MkP-restricted; T10, T cell-restricted. These ten clusters of clones were detected in recipients of donor E^hi^ HSCs from both unperturbed and 5-FU-treated mice (Figure 6E). However, compared to donor E^hi^ HSCs from unperturbed mice, those from 5-FU-treated mice produced more HSC-related clones (T2, T3, T6, T8) and fewer lymphoid-biased clones (T4) in the recipients (Figure 6E), suggesting increased self-renewal tendency and reduced lymphoid potential of E^hi^ HSCs following 5-FU treatment.

With the major differentiation patterns in the transplantation recipients determined, we assigned the 542 shared clones to one of the ten differentiation patterns and compared them to their differentiation patterns in the donors. First, a visual inspection found that most of these clones in the transplantation recipients showed differentiation patterns different from their differentiation behaviors in donors (Figure 6A). Indeed, the seven individual donor clone types produced most of the ten types of recipient clones (Figure 6F), suggesting that the differentiation patterns of these clones were not strictly inherited after transplantation. Furthermore, 123 of these 542 shared clones were detected in at least two recipients. Over two-thirds of these clones display diverse differentiation patterns in the individual recipients (Figure 6G), indicating the lack of differentiation imprinting in the clones.

In addition to the differentiation patterns, we also compared the overall self-renewal and differentiation activities of E^hi^ HSC in the 542 recurrent clones in donors and recipients. Among the 542 clones, we focused on the E^hi^/E^lo^/E^neg^ HSC and MkP unique donor clones and the C1, C2, and C4 multi-cell type donor clones, given their higher survival probability in the recipients. Among the shared clones derived from unperturbed mice, the C1 clones had the highest E^hi^ HSC UMI counts in the donor, as expected (Figure 6H), and the C4 clones had the highest combined UMI counts of HSC downstream cells (Figure 6I). This pattern was partially recapitulated in transplantation, with more recipient clones derived from the C1 donor clones showing high E^hi^ HSC UMI counts (Figure 6J) and more recipient clones of C4 donor clones showing high combined UMI counts of downstream cells (Figure 6K). Therefore, the high self-renewal potential of E^hi^ HSC in the C1 clones and the high multi-lineage differentiation potential of the E^hi^ HSC in the C4 clones were largely preserved after bone marrow transplantation. Consistent with the roles of HSPCs in 5-FU-induced hematopoietic regeneration, the clones from 5-FU treated mice showed higher levels of differentiation activity than those from untreated mice (Figure 6I). However, their differentiation activity was significantly reduced following bone marrow transplantation (Figure 6K), indicating prolonged defects induced by 5-FU treatment on the regeneration capacities of HSPCs in stress conditions (Hinge et al., 2020).

Together, these results indicate that selected HSC subsets partially preserve their self-renewal and multilineage differentiation abilities, but the exact differentiation patterns displayed in the unperturbed mice were not inherited under the transplantation condition.

## Discussion

Self-renewal and multi-lineage differentiation are hallmarks of hematopoietic stem cells (Eaves, 2015; Seita & Weissman, 2010). However, investigating these two properties in situ at a single-cell level has been challenging. As HSCs in adults primarily reside in a quiescent state and their progenies are mobile, it is difficult to capture the self-renewal and differentiation events of HSC using the imaging approach in live animals. As a result, various “barcoding” approaches have been developed to label the hematopoietic cells uniquely and irreversibly, and the differentiation behaviors are retrospectively read out after a certain length of chase, during which the initial barcodes are propagated into different cell types (Cordes, Wu, & Dunbar, 2021). In the current study, we used the Sleeping Beauty transposon system to label the hematopoietic cells at a young age unbiasedly and analyzed the self-renewal and differentiation behaviors in these clones after a lifelong chase. Over seventy thousand clones were analyzed in the current study. This large number of clones helps overcome the limitation in sampling and clone detection and facilitates the extraction of the general differentiation patterns in native hematopoiesis. The lifelong-chase period washes out most short-lived progenitors, so the clones analyzed at the end of the chase are primarily derived from HSCs.

Previous studies using limited dilution and single-cell transplantation approaches revealed balanced, myeloid-biased, lymphoid-biased, and megakaryocyte-restricted differentiation patterns in HSCs under the transplantation condition (Carrelha et al., 2018; Dykstra et al., 2007; Muller-Sieburg et al., 2004; Müller-Sieburg et al., 2002; Yamamoto et al., 2013). It is thus reassuring that similar types of differentiation bias were observed in unperturbed hematopoiesis. Moreover, detecting the HSPC populations in the same clones provides additional benefits to associating specific differentiation bias with the different self-renewal properties of HSCs. For instance, previous studies show that aging-related expansion of the HSC compartment is driven by clonal expansion of megakaryocyte-restricted HSC (Aksöz et al., 2024; Grover et al., 2016). Our results support this proposition by showing that the expansion of the HSC compartment, especially the most primitive E^hi^ HSC subset, correlates with the increased frequency of MkP-restricted clones. The high self-renewal tendency of HSCs in the MkP-restricted clones is also partially imprinted as they show higher levels of self-renewal upon bone marrow transplantation (Figure 6H, J). In contrast to the MkP-restricted clones, the E^hi^ HSCs from multipotent clones expand to a lesser degree in the HSC compartment but produce more downstream cells in both donor and transplant recipients (Figure 6H-K). These observations are consistent with the MkP-restricted HSC being the most primitive HSC subset (Carrelha et al., 2018) and further demonstrate that the self-renewal and multilineage differentiation biases are relatively stable traits in selected HSC subsets. In future studies, it will be interesting to investigate whether the MkP-restricted differentiation status directly promotes E^hi^ HSC self-renewal in addition to the mechanisms mediated by secretory factors of megakaryocytes (Bruns et al., 2014; Zhao et al., 2014).

Clonal hematopoiesis is a frequently observed phenomenon in the aged population (Miguel Ganuza et al., 2019; Jaiswal, 2020; Mitchell et al., 2022; van Zeventer et al., 2021). In humans, peripheral blood is often sampled to assess the levels of clonal expansion (Genovese et al., 2014; Jaiswal et al., 2014). As a result, it is challenging to determine the early differentiation steps in the bone marrow that lead to the clonal expansion in mature cells. In this regard, our data indicates that clonal expansion in different differentiated cell types is inherited from diverse progenitor cell types, even if all these clones in the aged mice likely originate from HSCs. For instance, clonal expansion in a subset of MkP is associated with the E^lo^ HSCs. In addition, myeloid expansion is closely related to the clonal expansion events in the HPC-1 or MyP compartments. These associations reveal the earliest steps in which the hematopoietic system expands to supply large quantities of mature cells.

Interestingly, although the clonal expansion in Gr and Mono can be traced back to the clonal expansion in HPC-1 and MyP, not all HPC-1 clones participate in the clonal expansion of downstream cells. By sophisticated clustering analysis, we discover differentiation-active and -indolent subsets of HPC-1 that are not correlated with specific lineage bias. This layer of heterogeneity in HPC-1 has not been previously reported. Such finding is due mainly to the higher diversity of Sleeping Beauty transposon, which makes it possible to find the common differentiation pattern from thousands of clones by unsupervised clustering. However, given the limitation in the design of the Sleeping Beauty transposon-labeling system, we cannot simultaneously obtain the transcriptional status of these clones. Thus, the molecular mechanism controlling these diverse differentiation behaviors remains to be determined.

Apart from unperturbed hematopoiesis, our study further investigates the impacts of myeloablative perturbation and bone marrow transplantation on the clonal behaviors of HSPCs. Consistent with the role of HSCs in 5-FU-induced hematopoietic regeneration, both MkP-restricted and multipotent HSC clones from 5-FU-treated mice produced more downstream cells during the lifelong chase than their counterparts from unperturbed mice (Figure 6I). However, this more active differentiation activity was not inherited under the transplantation condition (Figure 6K), consistent with the report that 5-FU-treatment compromises the engraftment ability of regenerated HSCs (Hinge et al., 2020). Therefore, 5-FU pre-exposure does not affect HSC differentiation in situ but does so under the transplantation condition.

Previous studies showed that the lineage bias of HSC differentiation is inheritable during rounds of transplantation. However, our analysis of unperturbed and transplantation hematopoiesis does not strongly support this conclusion (Figure 6F, G). We suspect that the limited number of clones simultaneously detected from donors and recipients, a prerequisite for comparing changes in differentiation bias, prevents a thorough evaluation of such a correlation. For instance, myeloid-biased clones cannot be easily recovered under the transplantation condition (Figure 6B, C), which precludes the analysis of these clones following bone marrow transplantation. On the other hand, we find that the self-renewal ability of HSCs in the MkP-restricted clones and the multilineage differentiation ability of HSCs in the multipotent clones are partially preserved following bone marrow transplantation (Figure 6H-K). This data indicates that the self-renewal and multi-lineage differentiation biases are relatively stable traits that can be observed under unperturbed and transplantation conditions. Uncovering how these traits are maintained at the molecular levels would strengthen our understanding of the mechanisms regulating HSC cell fates.

In summary, our study analyzes a large collection of hematopoietic clones to determine the connections between self-renewal and multilineage differentiation. These analyses reveal the fundamental properties of HSPC and confirm our previous understanding of HSC biology, such as HSC differentiation heterogeneity and the association between MkP-restricted differentiation and HSC self-renewal. Our study further demonstrates the distinct HSPC stages initiating clonal expansion in major blood cell types, the different HPC subsets with diverse engagement in cell differentiation, and the stability of self-renewal and multi-lineage differentiation traits in HSC subsets. These findings reveal HSC cell fates in their native bone marrow environment and pave the way for future studies uncovering HSC regulatory mechanisms.

## Material and Methods

### Mice

The M2/HSB/Tn mice were generated as previously described (J. Sun et al., 2014). The CD45.1 and CD45.2/CD45.1 mice were bred in-house. All mice were maintained on the C57BL/6J background. Both male and female mice were used in this study. Mice were randomly assigned to the different experimental groups. All mouse studies were carried out according to the guidelines of the Institutional Animal Care and Use Committee at ShanghaiTech University.

### Doxycyclin induction and 5-Fluorouracil treatment

To induce transposon mobilization in the M2/HSB/Tn mice for lifelong lineage tracing, 6-10-week-old mice were fed with 1 mg/ml Doxycyclin (Dox) and 5 mg/ml sucrose in drinking water for 4 weeks. To obtain granulocytes carrying the freshly mobilized transposon, Dox was dissolved in phosphate-buffered saline (PBS) at 10 mg/mL and injected retro-orbitally into 10-week-old M2/HSB/Tn mice at a dose of 25 mg/kg body weight 48 hours before blood collection. For 5-Fluorouracil (5-FU) treatment, 5-FU was dissolved in PBS at 8 mg/mL and injected intraperitoneally into mice at a dose of 150 mg/kg body weight. Mice were treated with three rounds of 5-FU three weeks after Dox withdrawal, with a 3-week interval between each 5-FU injection.

### Isolation of bone marrow cell populations

Bone marrow cells were collected by crushing bones, including femurs, tibias, ilia, and spine, in FACS buffer (2% fetal bovine serum in PBS) on the petri dish. After removing red blood cells (RBCs) with RBC lysis buffer, the bone marrow HSPC fraction was enriched using the CD117 microbeads (Miltenyi) and LS columns (Miltenyi). The cKit^-^ cell fraction was used to isolate mature cell types. Cells were resuspended in FACS buffer and stained with optimally titrated antibodies on ice. The following combinations of cell surface markers were used to define bone marrow cell populations, E^hi^ HSC: Lineage (Lin: Gr1, Ter119, B220, CD4, CD8a, CD11b)^−^ cKit^+^ Sca1^+^ CD48^−^ CD150^+^ EPCR^hi^; E^lo^ HSC: Lin^−^ cKit^+^ Sca1^+^ CD48^−^ CD150^+^ EPCR^lo^; E^neg^ HSC: Lin^−^ cKit^+^ Sca1^+^ CD48^−^ CD150^+^ EPCR^neg^; HPC-1: Lin^−^ cKit^+^ Sca1^+^ CD48^+^ CD150^−^; HPC-2: Lin^−^ cKit^+^ Sca1^+^ CD48^+^ CD150^+^; MyP: Lin^−^ cKit^+^ Sca1^−^ CD150^-^ CD41^-^; MkP: Lin^−^ cKit^+^ Sca1^−^ CD150^+^ CD41^+^; granulocytes: Ly6G^+^ Ly6C^-^ CD4^-^ CD8a^-^ B220^-^; monocytes: Ly6G^−^ Ly6C^+^ CD4^-^ CD8a^-^ B220^−^; B cells: B220^+^; T cells: CD4^+^ /CD8a^+^. DsRed^+^ fractions of the above cell populations were sorted to detect Tn tags. Cell sorting was performed on a FACS-Aria III flow cytometer (BD Biosciences). Gating strategies for cell sorting were shown in Figure 1—figure supplement 1, and antibodies used for staining were listed in Supplementary File 3. Flow cytometry data were analyzed using FlowJo software (BD Biosciences).

### Isolation of peripheral blood cell populations

Approximately three capillaries of peripheral blood were collected from the retro-orbital sinus. RBCs were removed by dextran-promoted aggregation and then by hypotonic lysis with RBC lysis buffer. After cell surface marker staining, the DsRed^+^ fractions of mature lineage cells were sorted as follows: granulocytes: Ly6G^+^ Ly6C^-^ CD4^-^ CD8a^-^ B220^-^; monocytes: Ly6G^−^ Ly6C^+^ CD4^-^ CD8a^-^ B220^−^; B cells: B220^+^; T cells: CD4^+^ /CD8a^+^. Cell sorting was performed on a FACS-Aria III flow cytometer (BD Biosciences). Gating strategies for cell sorting were shown in Figure 1—figure supplement 1B, and antibodies used for staining were listed in Supplementary file 3.

### Transplantation assays

DsRed^+^ E^hi^ HSCs from M2/HSB/Tn mice (CD45.2^+^) were transplanted through retro-orbital injection with 1.5 x 10^6^ whole bone marrow cells into lethally irradiated C57BL/6 recipient mice (CD45.1/CD45.2), which received two doses of 4.5 Gy with a four-hour interval. Cell sorting of peripheral blood cells was performed monthly for three months after transplantation. Bone marrow cell populations were sorted three months after transplantation.

### DNA isolation, Tn tag library construction, and sequencing

For Tn tag detection, transposase-assisted capture of transposable elements (TRACE) (Patel et al., 2022) was used with minor modifications. For samples with more than 100,000 cells, DNA was purified with GeneJET Whole Blood Genomic DNA Purification Mini Kit (Thermo Scientific). For samples with fewer than 20,000 cells, cells were incubated at 75°C for 30 minutes in 10 μL lysis buffer (60 mM Tris-HCl pH 8.0, 2 mM EDTA pH 8.0, 15 mM DTT) and then treated with 2 μL of 4 mg/mL Protease (QIAGEN) by incubation at 55°C for 6 hours, followed by 30 minutes of heat inactivation at 75°C. Otherwise, the volume of the lysis reaction of the sample was scaled up in equal parts in units of 20,000 cells. Genomic DNA from wild-type mice bone marrow granulocytes was spiked into samples with fewer than 20,000 cells to achieve a relatively equivalent total DNA amount of approximately 120 ng before fragmentation. Purified DNA or cell lysate was subjected to DNA fragmentation at 55°C for 12 minutes using an in-house prepared Tn5 transposase-adaptor complexes (Picelli et al., 2014) containing the Tn5MErev oligo and a customized UMI-bearing Tn5MEDS-BL oligonucleotide. The volume of Tn5 transposase used in each reaction was adjusted based on input DNA to ensure an average size of 400-500 bp for the fragmented DNA. The DNA-Tn5 mixture was purified with AxyPreP PCR Cleanup Kit (Axygen) to recover UMI-tagged DNA fragments. All of the purified DNA was subjected to a primary PCR amplification with 2×AceTaq Master Mix (Vazyme), the forward, sleeping beauty transposon-specific primer Tn3-1F, and the reverse Tn5MEDS-BL adaptor primer LCI. The Tn3-1F primer was biotinylated at the 5’ end, which allowed enrichment of the PCR products by using the G-Streptavidin Beads (PuriMag). PCR products were retrieved by alkaline denaturation using 5 μL of 0.1M NaOH at room temperature for 10 minutes, and 2 μL of the denatured product was used for secondary amplification with nested PCR primers MAF-N8-Tn3-1G and MAR-LCII, and further processed as previously described to generate libraries with 2 × Phanta Master Mix (Vazyme) for high-throughput sequencing on a NovaSeq PE150 platform (J. Sun et al., 2014). Tn tag identification and alignment were performed as previously described (J. Sun et al., 2014). The nucleotide sequences of primers used for Tn tag capture and sequencing library construction were listed in Supplementary file 4.

### Statistical analysis

To cluster clones detected in at least two cell populations based on the UMI counts of Tn tags, the UMI counts of each Tn tag in each cell population were first scaled as follows:

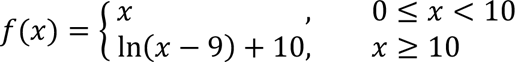

Subsequently, Louvain clustering was performed on the scaled UMI counts in the R package Seurat (Stuart et al., 2019). The number of neighbors was set to 50, and the resolution was manually adjusted to control the number of clusters.

To examine the level of clonal expansion in each population, Pielou’s index was calculated in each cell population with the following formulas:

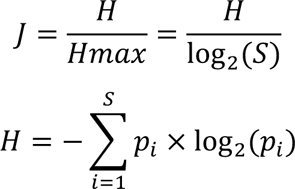

Where *J* is Pielou’s index, *H* is the observed diversity estimated by Shannon index, *S* is the number of observed clones, *Hmax* is the maximum diversity expected considering the number of observed clones (*S*), and *p_i_* is the frequency of UMI counts of clone *i* within the tested cell population.

To calculate the survival probability for each cell population, the number of Tn tags detected in both donors and recipients was divided by the number of Tn tags detected in donors.

To test whether HPC-1 clones could be classified into differentiation-active and differentiation-indolent groups, the Gaussian Mixture Model (GMM) analysis in the R package mclust (Scrucca et al., 2023) was performed on the clones in the C4 and C5 clusters based on the log2 ratio of UMI counts between upstream HPC-1 and downstream cell populations (MyP, Gr, or Mono). The log2 ratio was calculated as 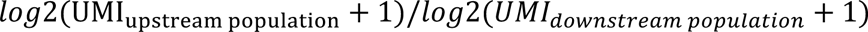. Clones that were not detected in HPC-1 were removed before analysis. The differentiation status of MyP to Gr and Mono is analyzed similarly.

To investigate the linear relationships of UMI counts between the cell population of interest and all of its possible upstream precursors, regression analysis was performed on their raw UMI counts using ElasticNet (Friedman et al., 2010; Simon et al., 2011) in the R package glmnet. The response family is set to “poisson (link=”log”)”. All other settings are set to default values.

To assess the statistical significance, we performed two-tailed unpaired Student’s t tests for two groups with GraphPad Prism 9 or Excel. A p value < 0.05 was considered statistically significant. n.s: not significant, *p < 0.05, **p < 0.01, ***p < 0.001.

## Data and code availability

The data files of Tn tags generated and analyzed during the current study, as well as the computer code used during the current study, are available from the corresponding authors upon reasonable request.

## Acknowledgment

The authors would like to thank Dr. Fernando Camargo for suggestions and sharing of the M2/HSB/Tn mice. We would like to thank the members of the Sun group for their discussion and support. We thank the Molecular and Cell Biology Core Facility (MCBCF) and Animal Core Facility at the School of Life Science and Technology, ShanghaiTech University, and the Animal Facility at the National Facility for Protein Science in Shanghai (NFPS), Zhangjiang Lab, Shanghai Advanced Research Institute, Chinese Academy of Science for providing technical service. We would like to thank the technical support of the HPC platform of ShanghaiTech University. This work was supported by the National Key R&D Program of China (2020YFA0710800, 2021YFA0804700 to JS) and the National Natural Science Foundation of China (81970102 to JS).

---

**Figure 1-figure supplement 1:**
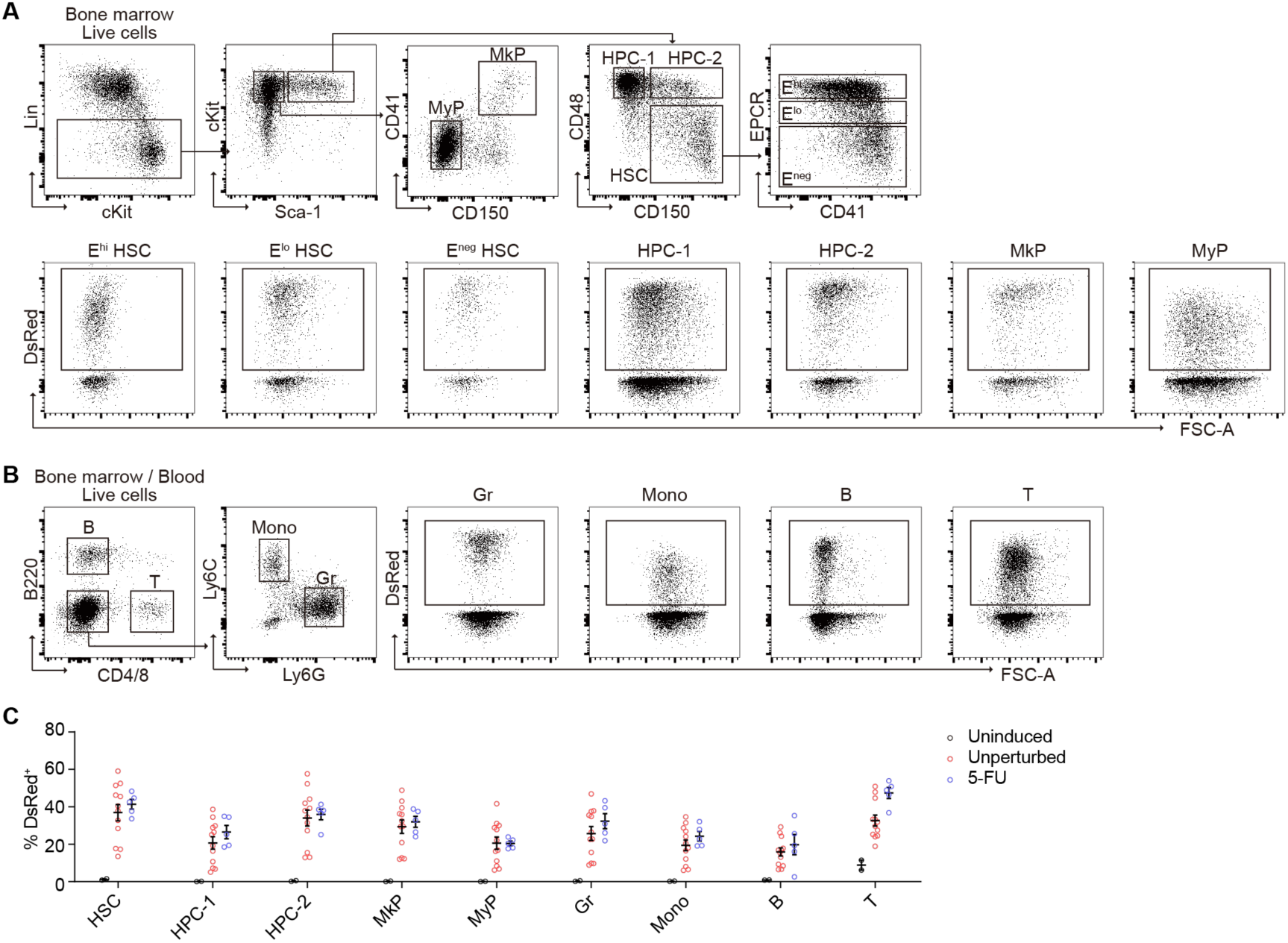
FACS gating strategy for cell isolation. A: The gating strategy used to identify and isolate E^hi/lo/neg^ HSC, HPC-1/-2, MkP, and MyP from cKit-enriched bone marrow cells. The DsRed^+^ fractions of the indicated cell populations are sorted for Tn tag detection. B: The gating strategy used to identify and isolate Gr, Mono, B cells, and T cells from cKit-depleted bone marrow cells or peripheral blood cells. The DsRed^+^ fractions of the indicated cell populations are sorted for Tn tag detection. C: Frequency of the DsRed^+^ fractions in the indicated bone marrow cell populations from aged mice sacrificed for Tn tag analysis. The two uninduced mice were analyzed at 120 and 123 weeks of age. Each circle represents data from an individual mouse (uninduced: black, n=2; unperturbed: red, n = 12; 5-FU: blue, n = 5). Bars represent means with SEM.

**Figure 1-figure supplement 2:**
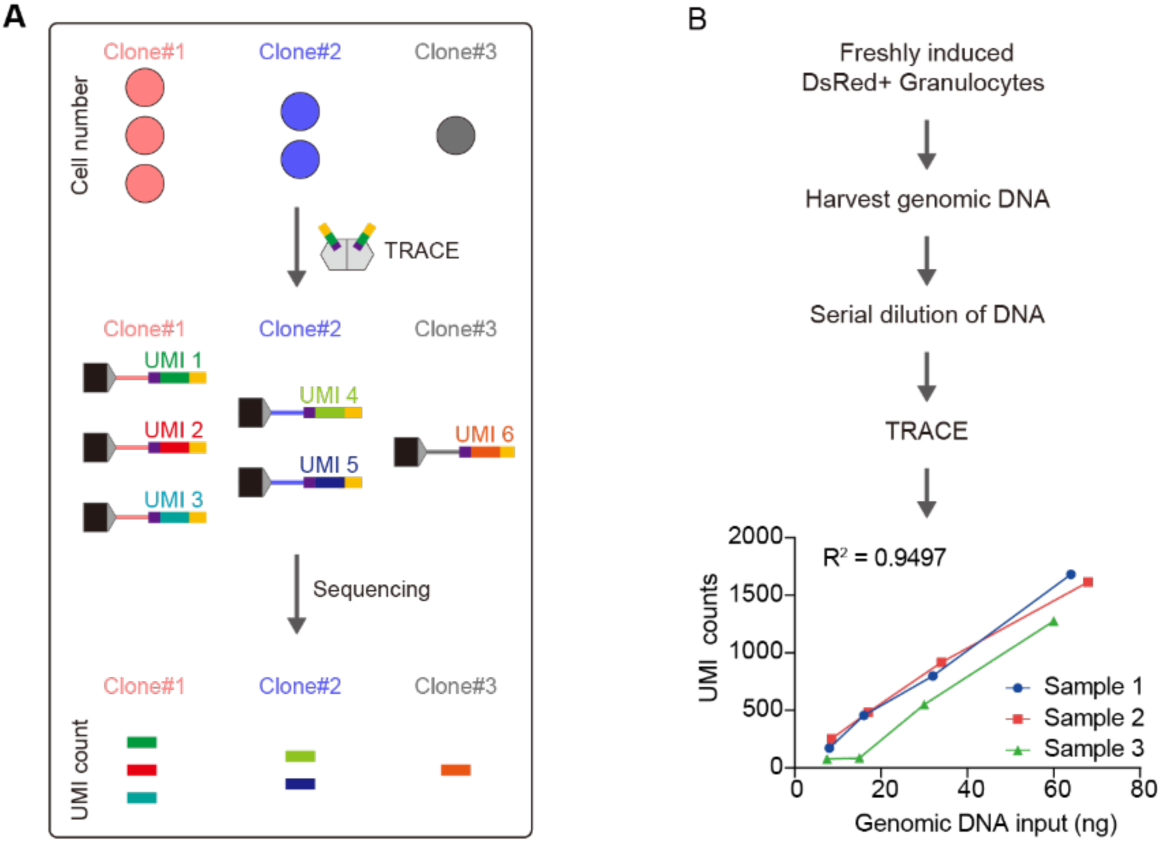
Characterization of the TRACE method for transposon tag detection. A: Schematic of the semi-quantitative TRACE method for Tn tag detection. B: Experimental design to validate the semi-quantitative property of the TRACE method for Tn tag detection. DNA from samples without clonal expansion (freshly induced granulocytes) is added at varying dilutions for Tn tag analysis. “R^2^” depicts the coefficient of determination of the linear regression.

**Figure 2-figure supplement 1:**
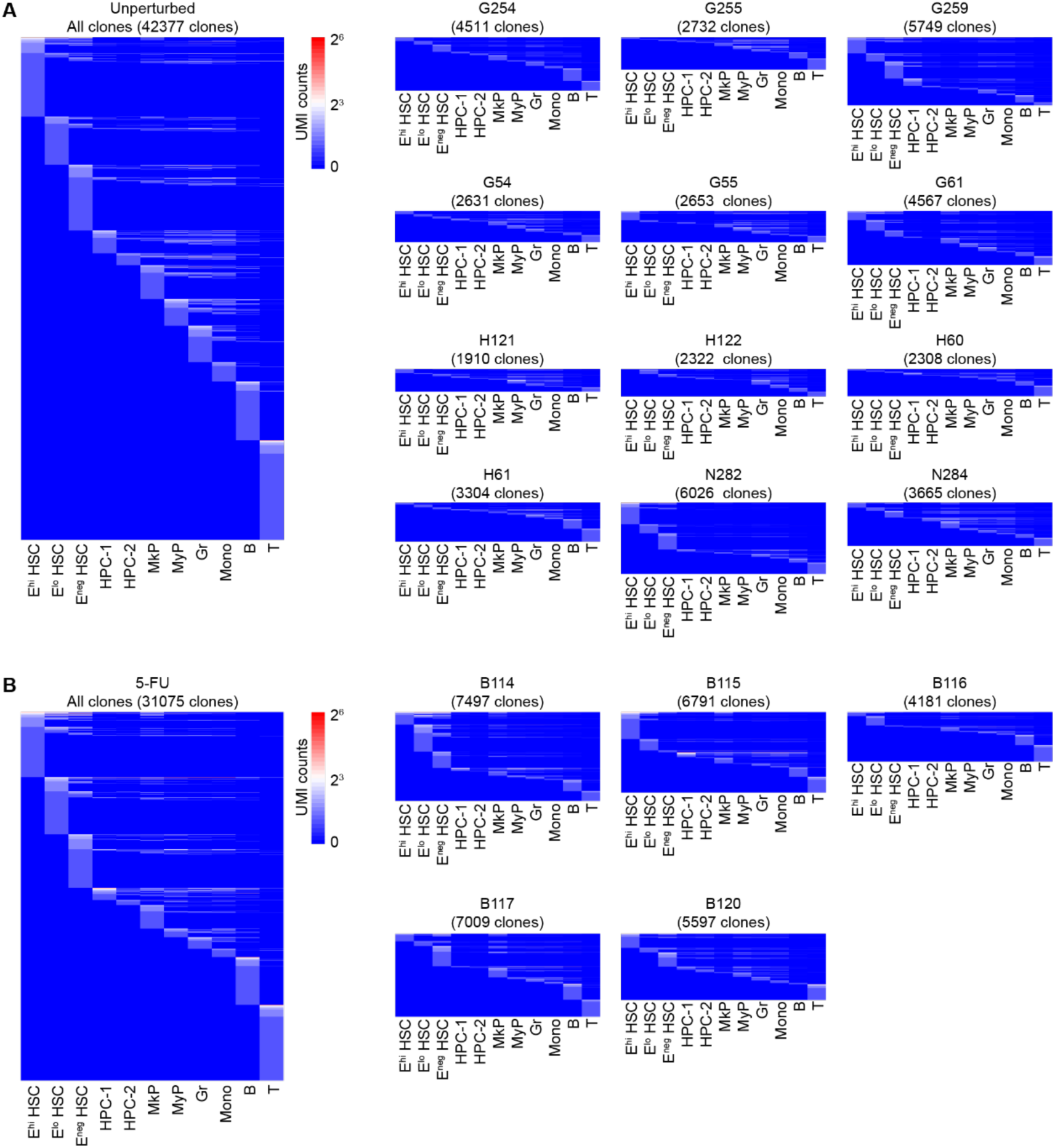
Alignment of transposon tags. A-B:Alignment of Tn tags across all assayed bone marrow cell populations, with each row representing a unique transposon tag and each column representing a sampled cell population. Tn tags are colored by their log2-transformed UMI counts, reflecting their abundance, and are arranged by rank. Tn tags detected in mice examined in the unperturbed (A, n = 12) and 5-FU-treated (B, n = 5) mice are shown togetheror individually.

**Figure 3-figure supplement 1:**
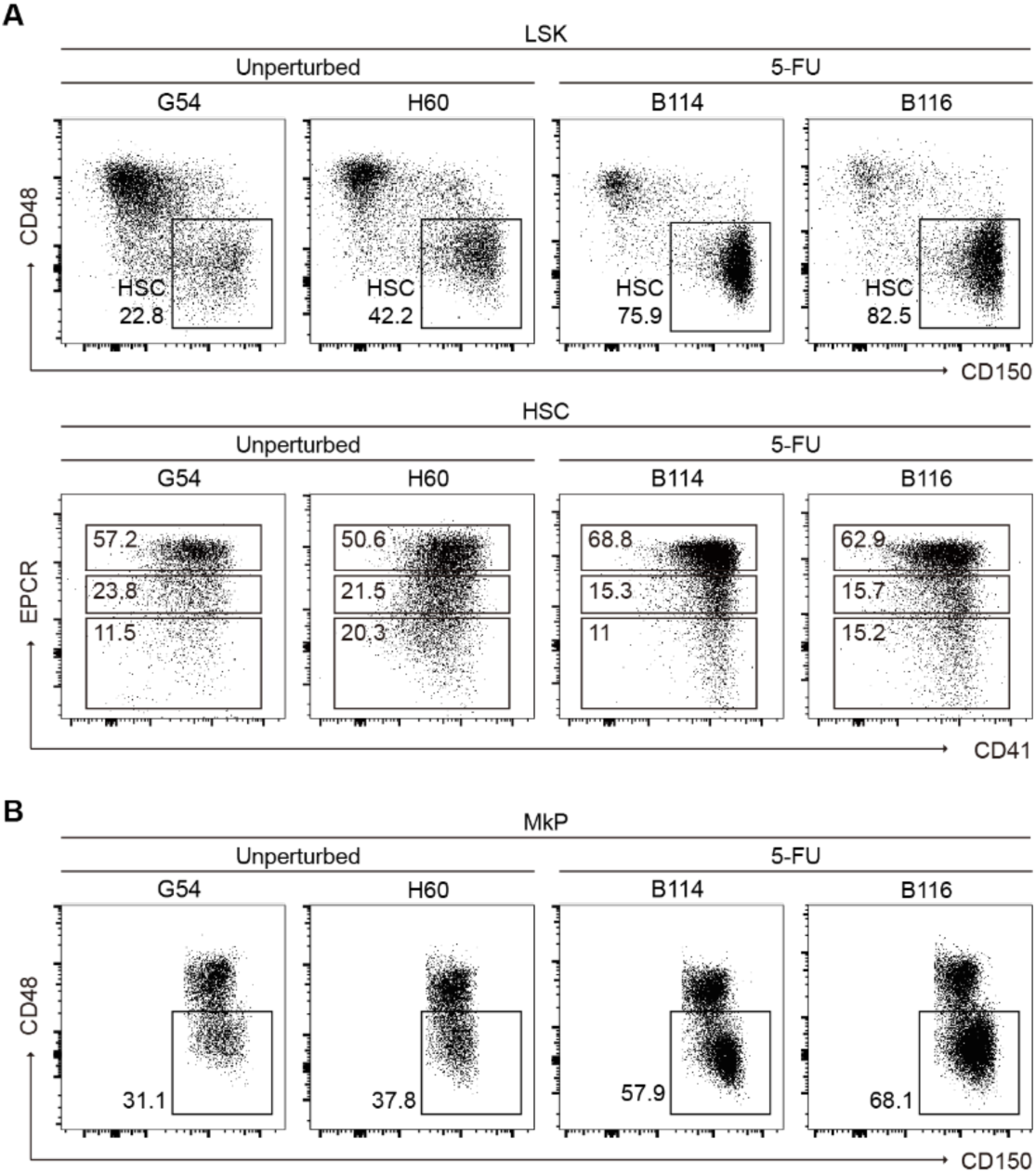
Expansion of HSC and CD48^-^ MkP after 5-FU-induced regeneration. A: Representative flow cytometry plots showing the frequency of HSC in Lin^-^ cKit^+^ Sca1^+^ (LSK) bone marrow cells from unperturbed and 5-FU-treated mice. B: Representative flow cytometry plots show the frequency of CD48^-^ cells in MkP from unperturbed and 5-FU-treated mice.

**Figure 4-figure supplement 1:**
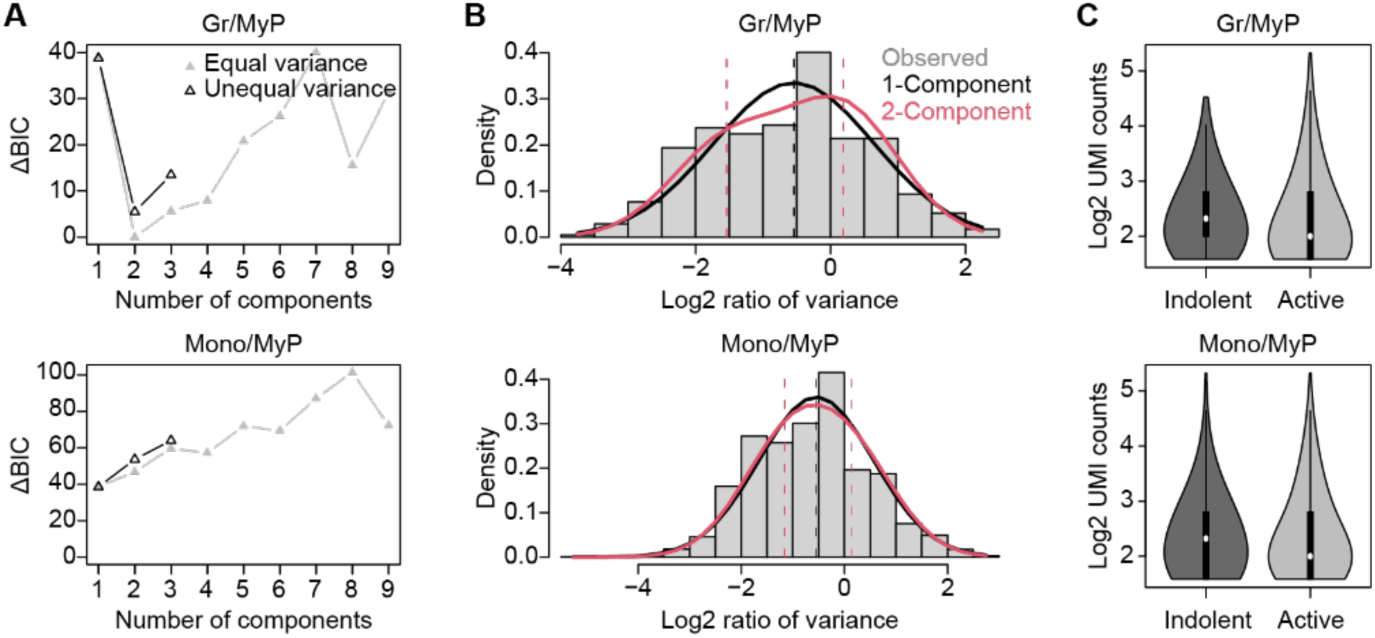
Gaussian mixture model classification of MyP by their differentiation states. A: The Δ Bayesian information criterion (ΔBIC) demonstrating the relative difference between the best model that explains the log2 ratios of UMI counts of Tn tags between downstream cell populations (Gr, Mono) and upstream MyP, and Gaussian mixture models with varying numbers of components. The results from models that assume each component has equal variance (gray) and unequal variance (black) are both shown. B: The observed distributions of the log2 ratios between downstream cell populations (Gr, Mono) and upstream MyP (gray bar) are shown alongside the distributions predicted by the one-(black curve) or two-component (red curve) Gaussian mixture models with minimal ΔBIC. The dashed lines illustrate the mean values predicted by the one-(black) or two-component (red) Gaussian mixture models. C: Violin plots showing the distribution of UMI counts of Tn tags detected in MyP in differentiation-active and -indolent clones. Data in this figure is pooled from mice assayed in the unperturbed state (n = 12) and after 5-FU-induced regeneration (n = 5).

**Figure 6-figure supplement 1:**
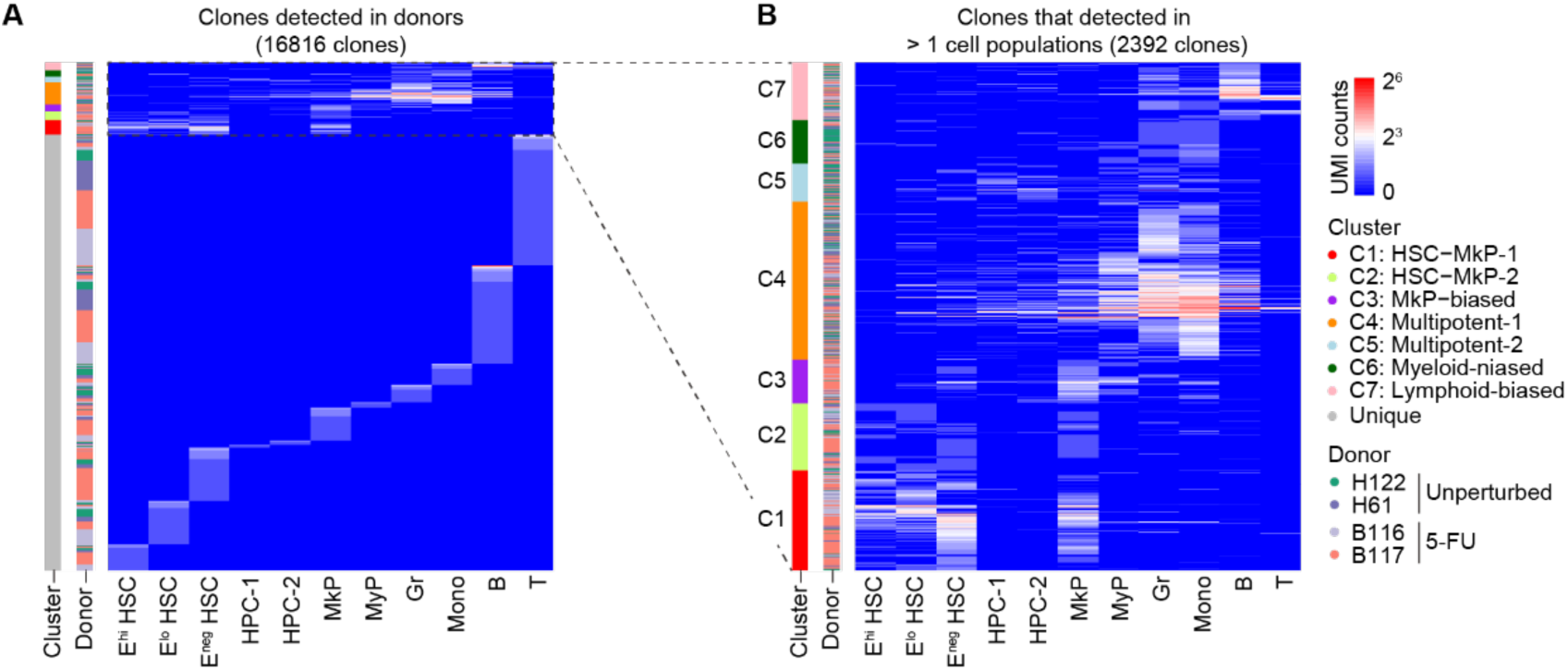
Alignment of transposon tags of donors. A-B:The heatmap showing all clones (A) and multi-cell type clones (B) detected in donor mice used for transplantation. Tn tags are colored by log2-transformed UMI counts. The transposon tags used in the plot are pooled from donor mice (n = 2 for each experimental group).

